# Macrophage-mediated epimorphosis in pig heart with right ventricular failure

**DOI:** 10.1101/2022.07.04.498652

**Authors:** V Lambert, A Deleris, F Tibourtine, V Fouilloux, A Martin, P Bridge, Bernd Jagla, MJ Arguel, L Gaigeard, P Barbry, E Aries, D Benoist, F Roubertie, M Pucéat

## Abstract

Heart left or right ventricular failure results from either ischemic or congenital diseases, respectively, and remains a major health burden in our societies. There is thus a high demand for a regenerative therapy. Yet, the ability of the adult post-mitotic mammalian heart to self-regenerate remains largely a challenge

Here, we combined cell therapy in a pig with right heart failure, cardiac physiology, single cell RNA-seq and spatial transcriptomics. We demonstrate that resident cardiac macrophages mediate a process of cardiomyocytes de-differentiation to form a blastema which produces new proliferative cardiomyocytes. Thus, a mammalian heart close to a human heart features the ability to undergo epimorphosis and to regenerate.

A direct and specific target of resident macrophages holds promise to regenerate hearts, specifically in a growing population of now adult congenital heart diseases patients with right ventricular failure and left without any efficient pharmacological relieving treatment.

---

Congenital heart diseases (CHD) are the major developmental diseases *(1)*. Significant and continuous progress in pediatric surgery from the 1990’s to repair cardiac malformations has greatly improved the survival of children who now constitutes a growing population of adult CHD patients. In this specific population, right ventricular (RV) dysfunction typically develops as a consequence of chronic pressure and/or volume overload which is secondary to the originally lifesaving surgical repair procedures *(2)*. Then, these young adult patients with grown up congenital heart (GUCH) diseases are at high risk of heart failure, severe arrhythmias, valve disease, pulmonary hypertension, cancer and potentially early aging *(3, 4)*, which constitutes a medical burden *(2, 5)*. Since more than 50% of GUCH patients have a risk of heart failure at 30 years old *(6)*, RV failure makes the long-term follow-up of CHD a challenging medical issue.

The RV is thin, features a gene profile different from the LV and a maladaptive metabolic response to afterload *(7)*, making it less equipped than LV to respond to overload. Furthermore, therapeutic options for this disease remain poorly investigated *(5)* and conventional therapeutic pharmacological approaches for these patients are often disappointing *(8)*. In the long run, if the repair of causal myocardial lesions does not allow for an improvement in function, or if RV is a single dominant ventricular heart such as in the hypoplastic left heart syndrome, organ transplantation is the only remaining option for these young patients. The difficulty and time to obtain heart grafts because of shortage of donors as well as the risk of such a heavy surgery for these young patients often severely affected *(9)* urgently calls for the search for alternative therapeutic pathways. Despite challenges which still need to be met, regenerative medicine still appears to be a powerful approach to prevent and treat terminal heart failure and its consequences *(10)*.

Clinical trials of cell therapy for congenital heart disease are still rare *(11, 12)*. The cell and more specifically the cell/molecular mechanisms as well as long term effects of that approach remained furthermore unclear.

The primary objective of our research, conducted on a large animal model of CHD *(13)*, was to elucidate the cellular and molecular mechanisms that underlie the effects of cell therapy.

To test the therapeutic effect of the graft of Cardiac Progenitor Cells (CPC), we performed heart surgery on 18 pigs in order to mimic the clinical situation of repaired TOF *(13)*. Four animals died during the study: three from the sham group (one from severe cardiac failure, two featured ventricular fibrillations), one from the control group (infection) before the end of the follow-up, and they were excluded from the final analysis. All remaining animals (n=6-8/group) gained weight and did not feature any clinical signs of heart failure. Age, weight, body length, and body surface area were similar in all groups at each stage (Fig. S1).

CPC were generated from the human embryonic stem cell line H9 genetically modified to express the ventricular myosin light chain 2 (MLC2v) fused to GFP *(14)* under the control of the cardiac α-actin promoter. Cells were sequentially challenged with a series of small molecules and growth factors (i.e, Wnt activator and BMP2 and then Wnt inhibitor and SHH) to direct them toward the fate of cardiac progenitors expressing NKX2.5 (Fig 1A) *(15)*. Cells were then sorted with an anti-CD15 antibody directed against an antigen (SSEA-1) expressed at the surface of differentiated cells *(16)* (Fig. 1A) to remove any undifferentiated cell as checked by the absence of OCT4+ cells (Fig S2A). CPCs were then seeded in a collagen/gelatin/alginate patch to cover the whole RV.

**Figure 1:**
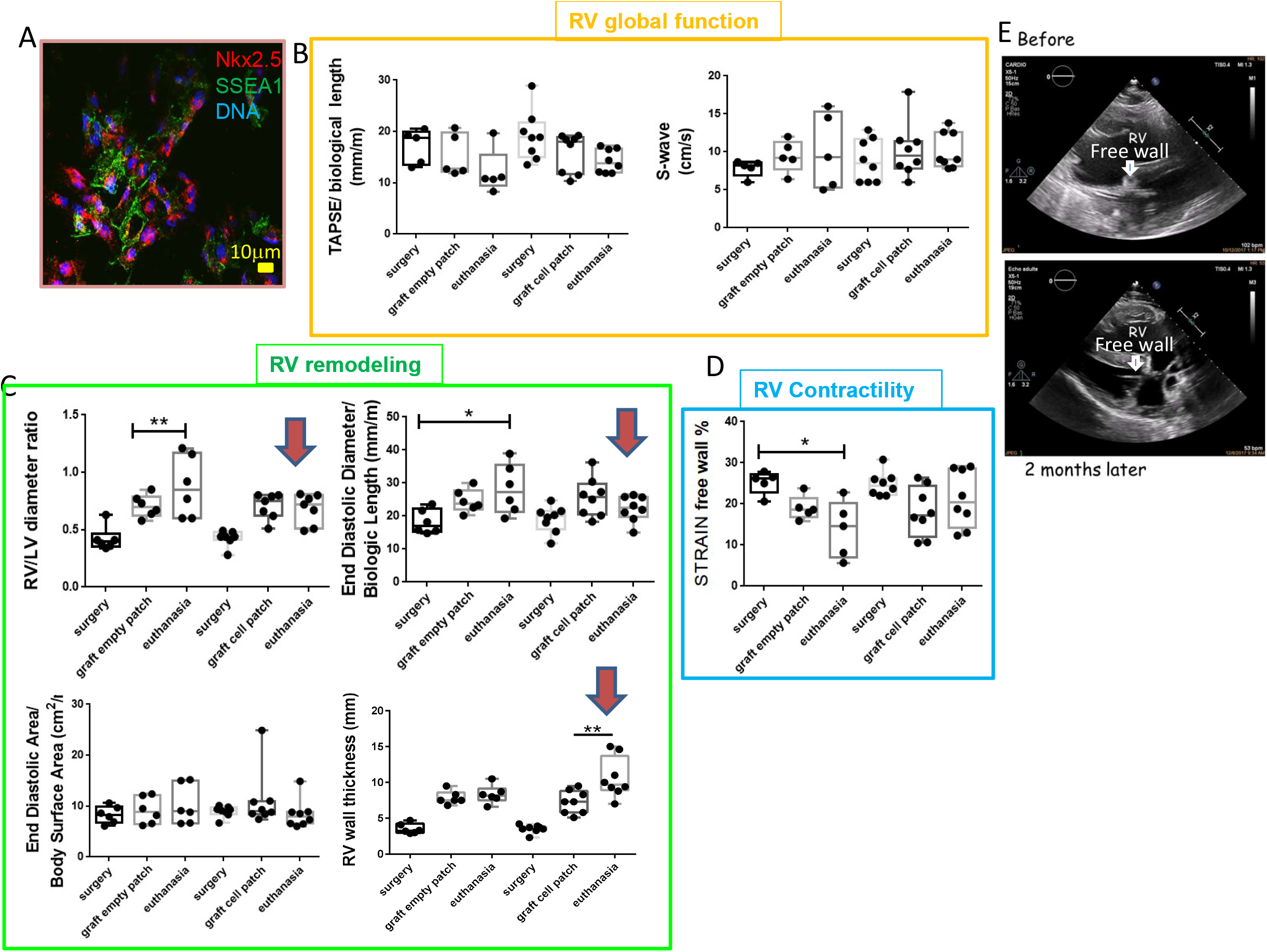
Functional parameters of pig grafted with cell-free patches. ** (empty) or (A) CPC–patch** (Nkx2.5+ cells, CPC expressing SSEA1) before the surgery (surgery), the day of the graft (graft) and at the end point of the experiment (euthanasia). **(B)** The top orange framed graphs represent the global function; **(C)** The left green-framed graphs reflect the remodeling of the RV. **(D)** the right blue framed, the contractility. T-test was used to compare two time points or Anova to compare 3 time points *p≤0.05 ** p ≤0.01 (**E**) Bidimensional echocardiography in a CPC treated animal: 4 cavities views, the right ventricle appears less dilated, the free wall thicker and the kinetic improved 2 months after cell pattch (down) compared to before (up) in the same animal.

The time between the surgery and the delivery of epicardial patches was similar between CPC seeded patch and cell-free groups (206.0 ± 24.8 *vs* 211.5 ± 40.7 days respectively) as well as the whole duration of the experiment (296.3±17.4 *vs* 292.8± 23.2 days respectively). No mortality was observed after cell delivery, or during the follow-up, and no adverse effects occurred. Immunosuppression was well tolerated. In the post cell therapy period, all animals gained weight (Fig S1B) and did not feature any clinical signs of either heart failure or immune rejection.

Five months after surgery, echocardiography revealed in the two groups a similar combined RV overload. A significant thickening of the RV free wall secondary to a moderate but significant pulmonary stenosis reflected a barometric overload, associated with a pronounced RV dilation (volumetric overload) assessed by increased RV/LV ratio, indexed diastolic diameter and both end-diastolic and end-systolic areas, following a severe pulmonary regurgitation (Fig.1). Echocardiographic standard parameters such as indexed TAPSE and peak systolic velocity of S’wave, as well as myocardial strain measures, significantly decreased to the same extent in the two groups (Fig 1B). These results showed an alteration of both global RV function and myocardial contractility. Two months after patch delivery, indexed RV diameter significantly decreased in the CPC group whereas it significantly increased in cell-free group. The free wall thickness significantly increased in the CPC group compared to cell-free group (Fig 1C,E) and the RV was less dilated than prior to the graft (Fig 1E). These results suggest that the RV adaptation to the overload was maintained and that a functional recovery took place. Furthermore, the myocardial contractility specifically assessed by free wall strain significantly improved in the CPC group in contrast to cell-free group in which it kept decreasing (Fig 1B and Fig S3). The free wall included an hypokinetic area which was no longer present after CPC-graft (Supplemental video data). Then, functional characteristics and contractility parameters differently evolved after either with or without cardiac progenitors (Fig 1D). Altogether, our findings show that RV function was no longer worsened in cell-grafted pig hearts and that the cell graft while limited could preserve the ventricular function.

Next, we searched for the cell and molecular mechanisms underlying the preservation of function in CPC-grafted pig hearts.

When grafted within the collagen-based patch covering the whole right ventricle of pigs, the human CPCs expressing phospho-histone 3 (PH3) still proliferated and were in the process of differentiation toward actinin+ cardiomyocytes within the patch two months after the graft (Fig 2A). However, most of cells migrated into the myocardium while differentiating into cardiomyocytes expressing the human nuclear antigen KU80 and fused MLC2vGFP protein within the sarcomeres (Fig 2B). Migration was likely accompanied with a degradation of fibrosis allowed by expression of metalloproteases (i.e. *MMP1* and *TIMP1)* by the CPCs (Fig. S2B). Interestingly, pig myocytes surrounding the human cardiomyocytes expressed NKX-2.5, a transcription factor not any longer expressed in adult myocytes (Fig 2A,E).

**Figure 2:**
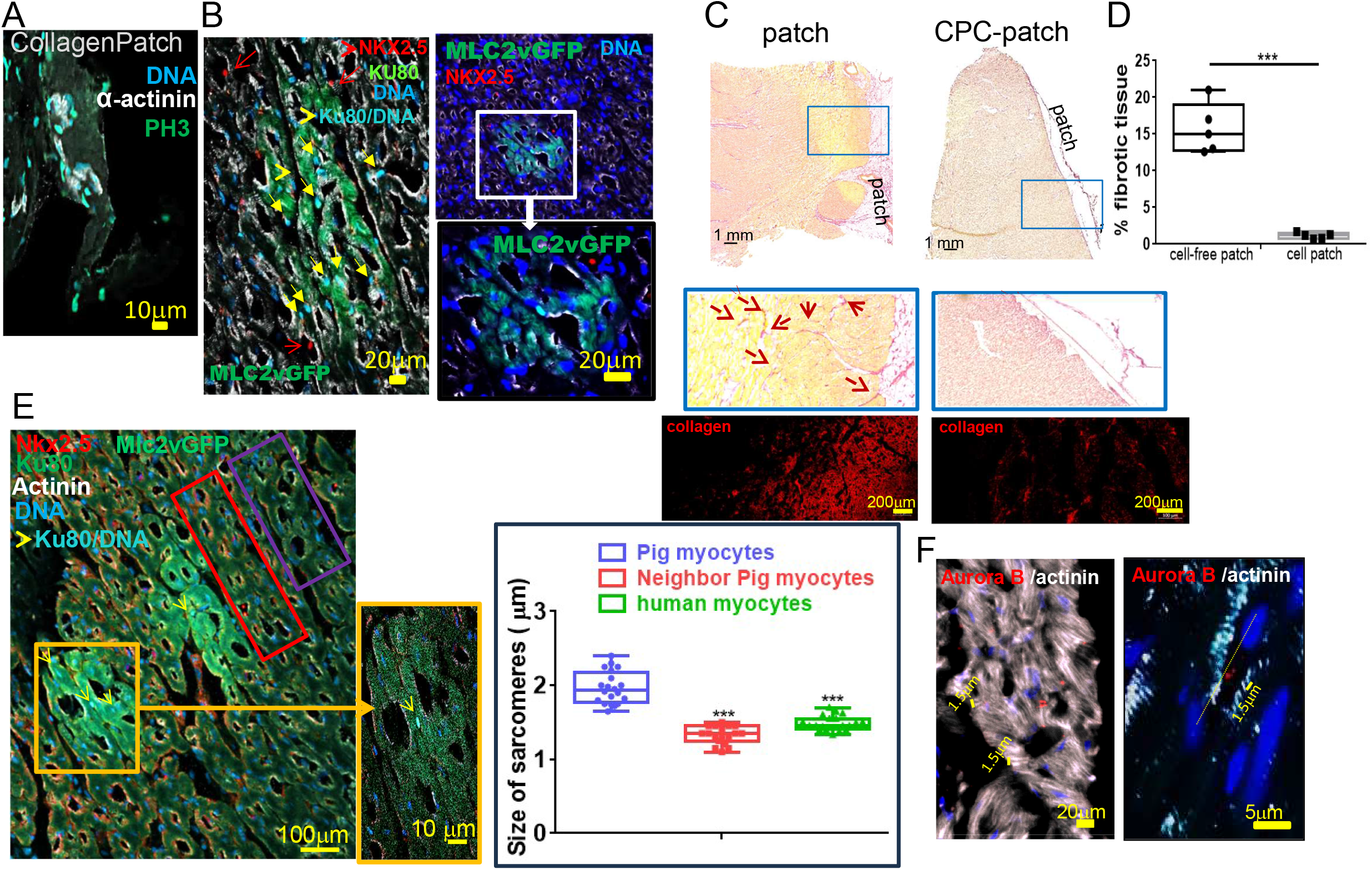
Human ES cell-derived Nkx2.5+ cardiac progenitors migrate, differentiate within the RV myocardium, trigger dedifferentiation and reproliferation of pig myocytes and induce regression of fibrosis. (**A**) human ES cell-derived Nkx2.5+ cardiac proliferate (PH3+) within the patch (**B**) migrate into and differentiate into clustered human KU80+ (light blue indicated by yellow arrow) MLC2VGFP+ cardiomyocytes. (**C**) Fibrosis is regressed in RV grafted with human cardiac progenitors. RV sections from pig grafted with an empty patch (Cell-free patch) or with a patch seeded with human cardiac progenitors (CPC-patch) stained with sirius red. Arrows indicate interstitial fibrosis. (**D**) quantification of fibrosis **(E)** size of sarcomeres measured in pig myocytes far or close to a graft of human myocytes in 3 separate hearts. (**F**) Dividing pig myocytes surrounding the graft were stained with an anti-auroraB antibody and sarcomeres lengths measured. The images are representative of sections from 5 different pigs **t-Test p<0.01

We next investigated the effect of the graft of human CPCs on the extent of fibrosis of the right ventricle. Pig hearts grafted with a cell-free patch still featured extended interstitial fibrosis as revealed by both Sirius red staining and monitoring of collagen fluorescence (Fig 2C). In contrast, pig hearts grafted with CPC-cellularized patches featured a restricted or no longer fibrosis (Fig 2C, D). Immunostaining with an anti-periostin to stain fibroblast or anti-smooth muscle actin antibody to stain myofibroblasts further confirmed a significant decrease in myofibroblasts in cell-grafted right ventricle (Fig S4A)

To further evaluate the reversal in the extent of fibrosis in human CPC-grafted right ventricle, we carried out RT-QPCR of genes expressed in fibroblasts or myofibroblasts. Expression of both *COL1A2* and *COL3A1* genes was significantly decreased in CPC-grafted hearts when compared to cell–free patch grafted right ventricle. Periostin *(POSTN)* expressed in fibroblasts and smooth muscle actin *(ACTA2)* expressed in myofibroblasts as well as the pro-fibrotic growth factor *TGFβ* were all downregulated in CPCs-grafted right ventricles (Fig S4B).

This intriguing extensive clearing of fibrosis led us to further investigate the phenotype of pig myocytes surrounding the human MLC2vGFP^+^ cardiomyocytes within the heart. Measurement of the size of sarcomeres revealed significantly shorter fetal like sarcomeres in pig myocytes neighbor of human cardiomyocytes than myocytes away from this area. Human myocytes also featured short fetal-like sarcomeres as expected from their still immature phenotype (Fig 2E). Aurora B staining and its specific localization in the midbodies showed that pig myocytes with short sarcomeres proliferated (Fig 2F). This was in agreement with the presence of clusters of early pig proliferating and differentiating myocytes negative for the human mitochondrial marker and which featured membrane troponin T (TnT) and nuclear Ki67 within the fibrotic area in the CPC-grafted hearts (Fig S5A). WGA staining revealed that pig cells with short sarcomeres1.43±0.035 μm were also smaller (52.7±4.5 μm), than adult pig myocytes (100± 5.6 μm, sarcomere of 2.38±0.043 μm) (Fig S5B).

As adult cardiomyocytes of 7 months pigs do not in principle reenter the cell cycle, we wondered how pig fetal like proliferating myocytes could have been generated around grafted human cardiomyocytes. We thus raised the hypothesis that pig myocytes could have de-differentiated and re-differentiated following reactivation of an embryonic genetic program. We thus looked for one of the major markers of pluripotent naïve stem cells, namely Oct4. We observed that some pig myocytes of CPC-grafted RV but not of cell-free patch grafted RV expressed OCT4 (Fig 3A). OCT4 was localized into the nucleus, thus likely to be transcriptionally active (Fig 3A, inset). Some cells also expressed NFκB or both nuclear OCT4 and peri-nuclear NFκB (Fig 3B). An anti-NANOG antibody failed to mark OCT4+ cells (data not shown) pointing to dedifferentiated and not reprogrammed cardiomyocytes. We thus reasoned that graft of CPC and their migration within the myocardium could trigger a local and spatially-restricted inflammation. Cytokines released within the environment could in principle activate NFκB pathway in either or both cardiac fibroblasts and cardiomyocytes. To test such a hypothesis, cardiac human fibroblasts were transfected with *p65/NF*κ*B* cDNA. Ten days later NFκB+ cells expressed OCT4 (Fig S6A). Second, neonatal post-mitotic day 7 mouse cardiomyocytes in primary culture were chosen for their capability to be transfected. *p65/NF*κ*B* cDNA was transfected in the myocytes. One week later, colonies of OCT4+ cells were observed in the culture (Fig S6B). These dedifferentiated cells re-differentiated into cardiomyocytes as indicated by re-expression of actinin set in still immature sarcomeres (Fig S6B). Interleukin 6 as well as oncostatin-M, two cytokines released by macrophages applied to myocytes also led to a dose dependent dedifferentiation in OCT4+ cells although cell dedifferentiation was not back to pluripotent stemness as NANOG was not expressed in these treated cells (data not shown).

**Figure 3:**
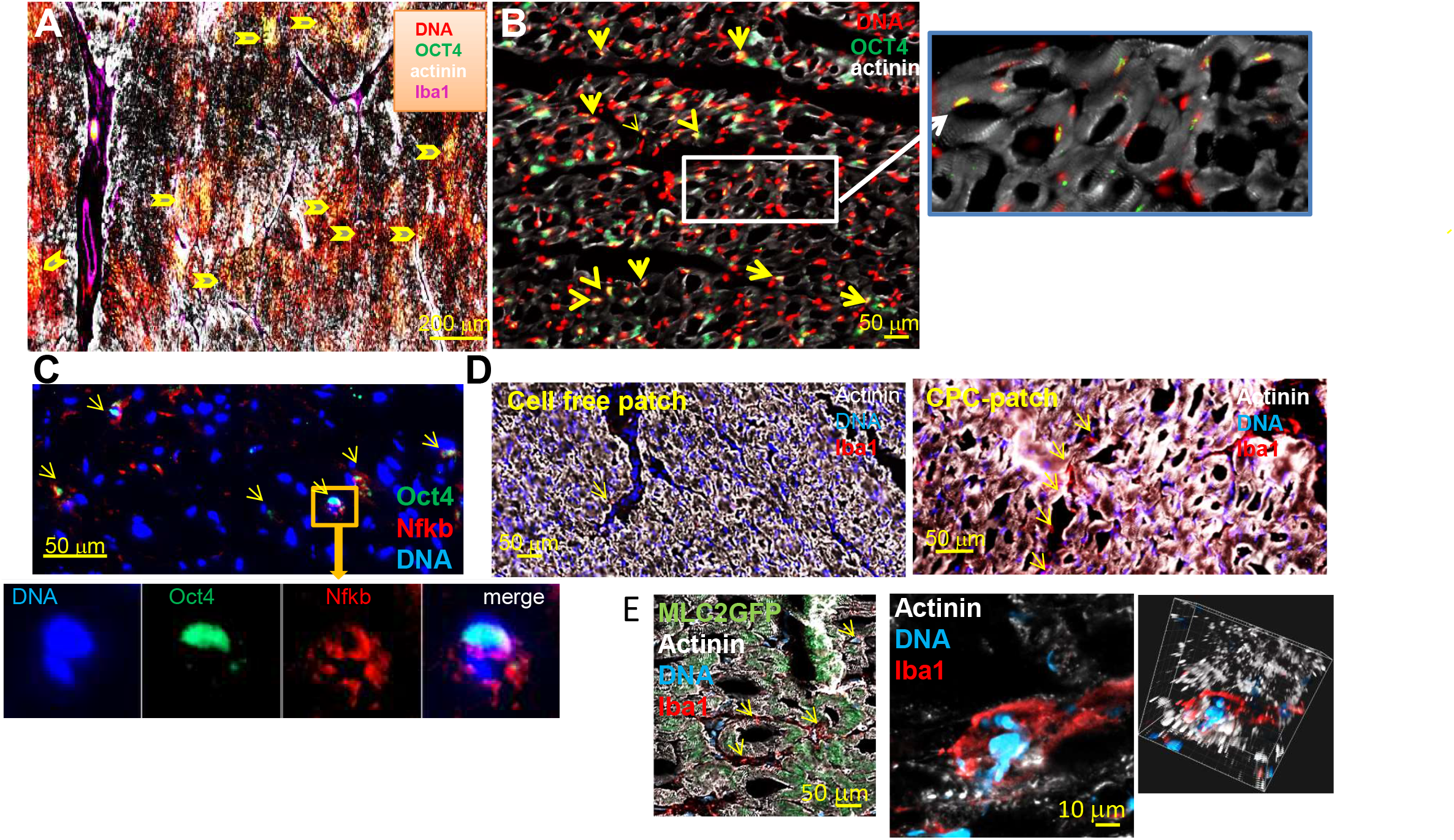
Dedifferentiated Pig myocytes reproliferate in human cardiac progenitor-grated RV. **(A)** clusters of Oct4+ cells in a large view of the CPC-grafted RV myocardial section also stained with anti-actinin (white) and anti-Iba1 antibody (purple). Arrows point oct4+ yellow cell nuclei. (**B**) a high magnification of cardiomyocytes from a CPC-grafted RV stained with anti-Oct4 antibody. The inset shows a higher magnification. (**C**) CPC-grafted RV sections were also stained with anti-Oct4 and anti-Nfkb antibodies. The inset shows a high magnification of Oct4+/Nfkb+ cells.The figures are representative of at least 4 different pigs grafted with cellularized patch. (**D**) Macrophages (Iba1+) staining in RV sections of pigs grafted with cell-free patch or CPC-patch. Yellow arrows point to macrophages. (**E**) Immunofluorescence anti-Iba1 on RV section of pigs grafted with CPC-patches within the graft area high magnification of Iba1+ macrophage on cardiac fibers. The right inset shows a 3D image reconstruction of the image revealing the position of the macrophage within the cardiac fiber.

We next hypothesized that a local inflammation should be mediated by resident immune cells. Macrophages have been involved in tissue regeneration *(17)* in many non-mammalian organisms. They are part of cardiac resident immune cells including in heart *(18)*. We thus tested the presence of macrophages in pig hearts. Iba1^+^ macrophages surrounded the OCT4^+^ cells in RV of CPC-grafted pigs (Fig 3D). While a few macrophages could be found in RV grafted with cell-free patches (Fig 3D), pig hearts grafted with CPC revealed the presence of many iba1^+^ macrophages and more specifically around the graft of human MLC2vGFP myocytes (Fig 3D). The immune cells were adherent to pig cardiomyocytes (Fig 3E) and 3D-reconstruction of a z-stack of images even showed that they were integrated within the cardiac fibers (inset Fig 3E) allowing for a close interaction and dialog with the myocytes. High CX3CR1^+^, Lyz2^+^ and TimD4^+^ macrophages from embryonic origin *(19)* were abundantly present two weeks after the graft (Figure S8).

To better learn about dedifferentiated pig myocytes in CPC-grafted hearts, single cells were enzymatically isolated from right ventricle two weeks after the graft of human CPCs. More specifically sus-epicardial cells were isolated and subjected to single-cell RNA-seq. The Heatmap (Fig.S9), the Uniform Manifold Approximation and Projection (UMAP) (Fig. S9B) as well as the panel plots (Fig. S9C) revealed the different cell types included in the whole population of isolated cells. The endothelial/endocardial cells were the most represented cell types as expected in adult heart *(20)* (Fig S9A,B). These cells were composed of both proliferative (cluster 0) likely sprouting endothelial cells indicating a remodeling of vessels required for tissue regeneration *(21)* and quiescent p21^+^ *(CDKN1A+)* cells (cluster1 and 7) as expected in adult organ. We also found *MSX1*^*+*^ cells undergoing endothelial-to-mesenchymal transition (cluster 4). More interestingly, we found a population of highly immature cardiomyocytes still expressing cardiac transcription factors such as *NKX2-5, GATA4, MEF2C* as well as *ACTN2* and very early cardiac genes such as *BVES, FHOD3, TNNI3 (22)* (cluster 6) no more expressed in mature cardiomyocytes (cluster 2) and whose expression was not detected in non-grafted RV pigs (Fig. S9). Besides smooth muscle cells (cluster 8) and ATP-binding cassette sub-family member 9 *(ABBC9*+) pericytes (cluster 3), we also identified cells expressing fibroblastic markers *PDGFRα, SERPINF1, COL3A1, COL1A1*, and high level of lumican *(LUM)*, the regenerative extracellular matrix *TNC, DPT* genes as well *LGI2* suggesting stimulation of nerves (cluster 5, Fig S9A,B) as previously observed in regenerative situation *(23)*. All these genes are highly expressed in blastema cells of axolotl *(24)* as well as in mammalian ear-hole in the course of epimorphosis *(25)*. Two populations of macrophages *(18)* including *LYVE1+, CD163+, CCR2*- and oncostatin+ (OSM) resident cardiac macrophages and *AIF1+, PLEH1+, TYROBP+* were also present (Fig S9A,B,C) and were sub-clustered in clusters 9 and 10 (Fig. S9D). Interestingly the highly immature cardiomyocytes (cluster 6) (Fig S9E) were the only cells to express many genes reminiscent of glycolysis, Krebs cycle and a few involved in fatty acid metabolism (Fig S9F), suggesting a high metabolic glycolytic demand as expected in fetal and proliferative myocytes *(26)*. This metabolic status also facilitates cardiac regeneration *(27)*.

To gain further insight into the dedifferentiation process, we used a spatial transcriptomics approach. Cryosections of CPC-grafted and empty patch grafted RVs were analyzed two weeks after the graft by spatial transcriptomics on a Xenium device. A pig specific (sus-scrofa) panel of 218 (ID 7HUPAE) genes (supplemental data) was designed (genome Sscrofa 11.1) to reflect the different cell types found in our single cell-RNA-seq experiments (Fig. 4). Using this set of genes, we could observe the main cell types in RV from control or CPC-grafted patch. Cardiac progenitors surrounded by blastema cells were only present in CPC-grafted RV. The cardiac sections looked less fibrotic and more cellularized than control RV (Fig 4, S11).

**Figure 4:**
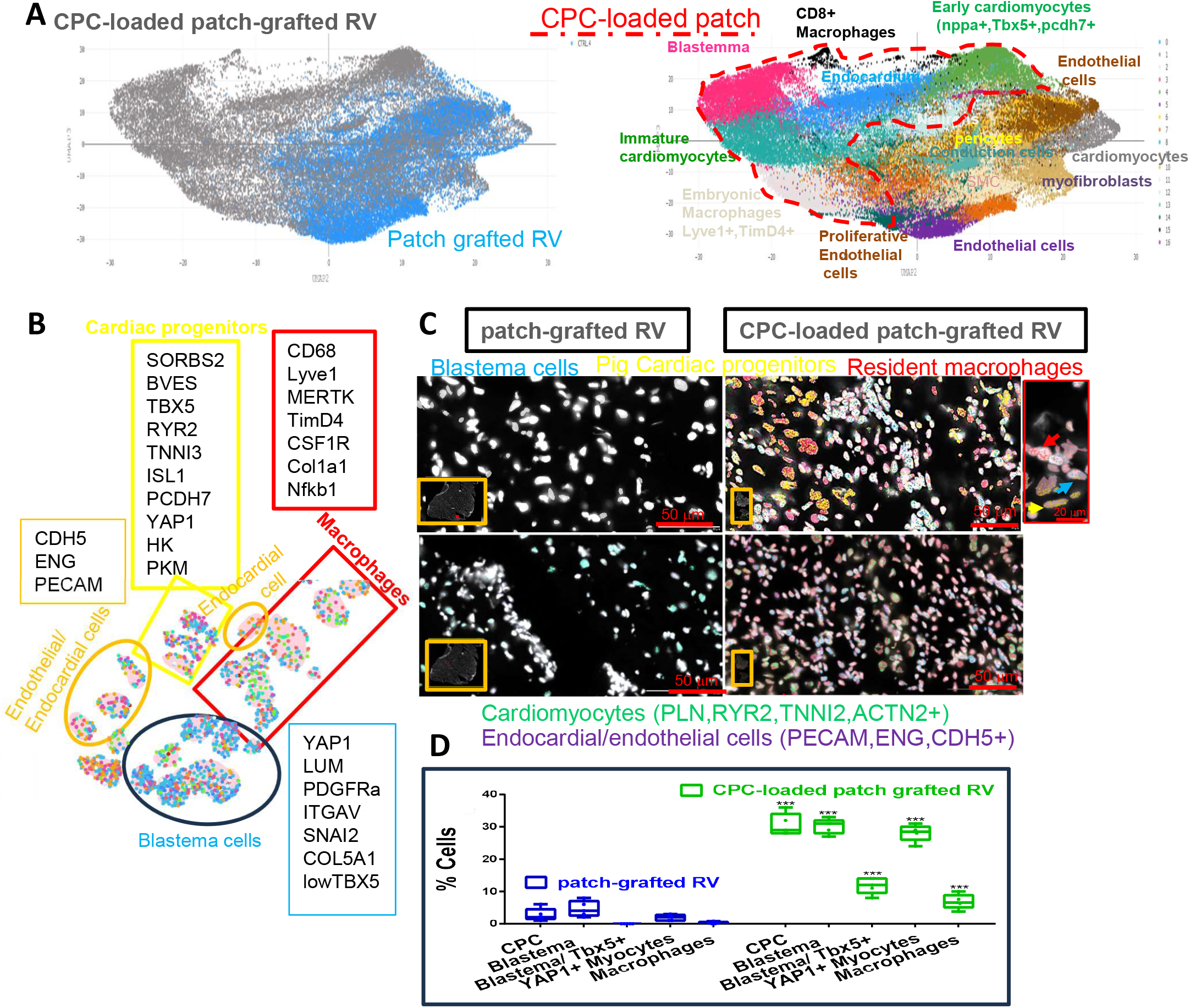
Spatial transcriptomics from pig heart slices grafted with CPC-loaded or empty patch. **(A)** UMAPS from 500 000 cells of each sample (2 combined control from empty patch and 2 combined CPC-loaded patch (B) a ROI on a slice of CPC-loaded patch RV showing specificity of cell types and genes used to identify them (C) Areas from Xenium-generated images from RV sections from slices grafted with CPC-loaded or empty patch showing different cell types present in these sections; right inset shows a magnification of cells to visualize nuclear transcripts (D) cell types scoring in 5 areas from 2 sections of VDs.

UMAPs in Figure 4A provided more insight into different cell types present in CPC-grafted RV than in in empty patch grafted RV (control RV). More specifically, CPC-loaded patch grafted RV were characterized by clusters of cardiac *TBX5*^*+*^, *ISL1*^*+*^, *BVES*^*+*^, *SORBS2*^+^, *RYR2*^*+*^, *TNNI3*^*+*^, *PCDH7*^*+*^, and *YAP*^+^ dedifferentiated cells expressing glycolysis genes *HK, PFM*, of dedifferentiated blastema cells expressing *LUM, PDGFRA, ITGAV, SNAI2, COL5A1*, as well *LLRC5, MSX1* and low *TBX5* (i.e. an index of their dedifferentiation status) as well as an over representation of resident macrophages. (Fig 4A,B,C S11) in CPC-grafted RV than in control RV in which CPC, blastema and macrophages were barely detected (Fig 4D). Interestingly these cell types were all neighbors when observed in a RV slice including cardiomyocytes and many endocardial/endothelial cells (Fig 4C,D S11).

Here, we demonstrate that the epicardial delivery of cardiac progenitors in pigs with right ventricular failure activates repairing macrophages and triggers dedifferentiation of pig cardiomyocytes into OCT4+ cells featuring a transcriptomic signature of blastema cells. The later re-differentiate into ventricular myocytes possibly with a left ventricular identity (TBX5+). This regenerates the myocardium. The cells allow fibrosis to be reversed and improve index of contractility of the ventricle following a true myocardial regeneration process (Fig S12).

The pigs used in this study were 6-7 months old at the time of cell grafting thus far from the regenerative post-birth windows *(28, 29)*. Furthermore, the *in-vivo* myocyte de-differentiation was triggered by a cell-cell interactions and endogenous cytokines without delivery of any pluripotency genes previously reported to trigger the same process s*(30)*.

The role of inflammation and specifically macrophages in tissue regeneration is not entirely new *(31, 32)* but the cell and molecular mechanisms have not been well documented, specifically for myocardial regeneration. It has furthermore never been directly associated with a cell therapy approach or involved in a process of *in vivo* cell de-differentiation.

We now show that the primary event and the trigger of myocardial regeneration is the migration of cardiac progenitor cells from the epicardial patch towards the myocardium. The cells which underwent epithelial-to-mesenchymal transition earlier in the process of differentiation, an event required for their migration recapitulating the embryonic scenario indeed have the proteolytic equipment (Fig S2) to degrade the proteins of the extracellular matrix. They thus make their own path toward the myocardium through the interstitial fibrosis while differentiating (Fig 1). The degraded proteins thus add to the damaged associated molecular patterns (DAMP) likely constituted of components of dead myocytes or released by them such as ATP. This triggers an inflammatory process as revealed by the presence of NFκB expressing cells (Fig 3, Fig 4) still present two months after cell delivery. Interestingly many cells including fibroblasts and cardiomyocytes feature DAMP-sensing receptors *(33)* making the scenario more than likely.

The activated NFκB pathway in cardiomyocytes triggers a re-expression of the pioneer factor Oct4 as revealed by immunofluorescence of CPC-grafted right ventricular sections (Fig 3). That NFκB is sufficient to induce Oct4 and to reprogram both cardiac fibroblasts and myocytes was revealed by our in vitro cell transfection assays (Fig S6) and as we previously reported for valvular cells *(34)*. These Oct4^+^ cells look similar to blastema cells (Fig 3) found in regenerative species such as amphibians *(35)*.

Interestingly, the Oct4+ cells are capable to re-differentiate into myocytes both *in-vivo* and *in-vitro* (Fig 3 and S5) likely because of their epigenetic memory that they retained as a print of blastema cells (26).

The immune cell involved is likely to be a macrophage often reported as a pro-regenerative and tissue repair cell. We found tow populations of macrophages in our single cell-RNA seq and spatial transcriptomics analysis (Fig. S8, 4). Iba1+ macrophages were also attached to cardiomyocytes and clustered in RV grafted with CPC around the graft, which suggests their activated state (Fig 3). They thus have a privileged location to interact and dialog with cardiomyocytes and in turn to secrete their cytokines in their vicinity. Which macrophage plays that role is still not yet known under our experimental situation. We can however surmise that it could be CCR2^-^, LYVE1^+^, CD163^+^, TIMD4^+^ macrophages of embryonic origin known to play a role in pediatric heart regeneration *(18, 19, 32)*. Lavine et al *(32)* have reported that embryonic macrophages are replaced by pro-inflammatory monocytes in adult post myocardial infarcted (MI) mouse hearts although adult heart retained a pool of self-renewing macrophages of embryonic origin *(36)*. However, our pig model of RV failure with interstitial fibrosis is different from a mouse MI model with a scar. Furthermore, Pig macrophages are transcriptionally closer to human than mouse immune cells *(37)*. Local activation of CCR2^-^, LYVE1^+^, CD163^+^, TIMD4^+^macrophages within the interstitial fibrosis is still a very likely scenario. Interestingly, activation of macrophages could help in clearing the fibrosis by a facilitated infiltration within proliferating new cardiac myocytes *(38)*. Macrophage activation and efferocytosis also induce macrophage proliferation to further resolve tissue injury *(39)*. Such a proliferative process for new cardiac progenitors emerging from a blastema could also be boosted by many p21^+^ quiescent or senescent cells that we identified in our single cell-RNA seq dataset (Fig 4) as found during axolotl regeneration process *(40)*.

How NFκB and cardiac inflammasome do dedifferentiate pig myocytes without triggering an adverse inflammatory process remains to be elucidated. The location and the intimate dialog of specific resident embryonic macrophages with myocytes could in part answer the question. Furthermore, NFκB acts as a pioneer factor and likely requires another transcription factor to bind super-enhancer on *OCT4* gene regulatory regions *(41)*. It acts as a dosage dependent manner and its target-genes depend upon the frequency and amplitude of its intracellular oscillations *(41, 42)*. Thus, a time-dependent fine-tuning regulation of activation of NFκB likely allows for cell reprogramming *versus* inflammation and fibrosis.

Microglia (i.e macrophages) that are also from embryonic origin *(43)* have been recently reported to promote adult brain repair as well as liver regeneration, an event mediated by an IL6 pro-regenerative pathway *(44, 45)* even if the mechanism of the regeneration was not investigated.

Dedifferentiation or even reverse differentiation into blastema cells as reported herein could be an efficient means to organ regeneration as previously reported and debated *(46, 47)*. Oncostatin has been suggested to dedifferentiate cardiomyocytes *(48, 49)* as we found (Fig

S7). Furthermore, the cytokine inhibits TGF*β*-dependent fibrosis *(50)*. Dedifferentiation or even reverse differentiation is as more important and even facilitated for organs such as the hearts that lack stem cells, which could compete with such a process *(46)*. Thus, such a macrophage-induced cell reprogramming or reverse differentiation toward a blastema-like stage as still functional for mammalian finger tips *(51)* could be a general cell mechanism to be explored in innovative regenerative medicine.

## Supporting information

echocardiography before the graft

echocardiography 2 months after CPC graftthe graft

## ACKNOWLEGMENTS

We are very grateful to de Federation Francaise de Cardiologie (To VL, MP) and The Leducq Foundation which funded this work (SHAPEHEART network of excellence) and for generously awarding us for cell imaging facility (MP “Equipement de Recherche et Plateformes Technologiques” (ERPT)). We thank Drs Serge Van de Pavert and Marc Bajenoff (Centre Immunologie de Marseille-Luminy) for fruitful and helpful discussions. We also thank Dr Stéphanie Chuun (Necker hospital, Paris) for screening the tacrolimus concentration from pigs blood. We thank for their support UCA GenomiX from the National Infrastructure France Génomique [ANR-10-INBS-09-03 and ANR-10-INBS-09-02], the 3IA@ coted’azur [ANR-19-P3IA-0002], the PPIA 4D-OMICS [ANR-21-ESRE-0052], the Conseil Départemental des Alpes Maritimes (2016-294DGADSH-CV) and the PEPR CellID [ANR-24-EXCI-0002].

## MATERIALS AND METHODS

### Genetic modification of HUES cells and cardiac differentiation

H9 HUES cell line was electroporated with a human α-actin pEGFP-C2 MLC2v DNA construct*(14)* and the clones selected using 100 μg/ml geneticin for 10 days. Cells were differentiated as previously described*(15)*. Briefly, cells cultured on MEF were stimulated with 10 μM ChIR9901 for 14hrs (overnight) and then for 24 hrs with 5 μM ChIR9901 and 10 ng/ml BMP2, in RPMI medium supplemented with B27. The cells were further incubated with IWR1 10 μM and BMP2 for 24hrs and then with 2.5 μM IWP4 with 10 ng/ml BMP2 and finally for one more day with 2.5 μM IWP4 and 10 ng/ml SHH. Cells were then sorted using magnetic beads conjugated with an anti-CD15 antibody (clone MC-480) (Kit easy step, Stem cell Technology).

### Cell isolation, differentiation and transfection

Human cardiac fibroblasts were from Promocell, (France). Neonatal mouse cardiomyocytes were isolated from 6 days old mice as previously reported *(52)* and isolated on a Percoll gradient.

Both cell types were transfected in Optimem (ThermoFisher, France) using Trans IT (Mirus Bio) according to the manufacturer protocol. Human cardiomyocytes were differentiated from H9 TnTGFP embryonic stem cells using CHIR9921 (MedChemExpress, Sweden), 5 μM for 2 days with 10 ng/ml BMP2 (MedChemExpress, Sweden), added the second day in RMPI-B27 medium, 5 μM XAV 939 (MedChemExpress, Sweden), for the next two 2 days and then only the medium for 5 days.

### Pig surgery and cell patch setting

All experiments were carried out according to the European Community guiding principles in a surgical procedure mimicking repaired tetralogy of Fallot (rTOF) was performed on 18 Landrace male piglets of two months old to induce overloaded RV dysfunction. Briefly, an enlargement of the RV outflow tract by a polytetrafluoroethylene patch, excision of one pulmonary valve leaflet, and a pulmonary artery banding were performed through a left thoracotomy approach*(13)*. After 5 months (median: 145 ± 22 days) of a combined pressure and volume overloaded RV which leads to its contractile failure. A large patch composed of collagen mixed with 10^7^ cardiac Nkx2.5+ progenitors (CPC), gelatin and alginate polymerized with 100 mM Ca2+ was sunk in a mold with the size of the RV. The patch was surgically set on the epicardium through a right thoracotomy approach, to wrap the whole RV free wall of 8 pigs (CPC group). A control group (n=6) included pigs which received a cell-free patch. These patches are scalable and their stiffness (0.1 kPa) allowed an easy handling for the surgeons. The patch remains stuck to the epicardium without any suture after closure of the pericardium. Hydrocortisone (1mg/kg) was injected intravenously before closing to reduce inflammation. All animals were immunosuppressed by Tacrolimus (daily 0.3 mg/kg of Advagraft ®) from 10 days before cell therapy until euthanasia performed after a median delay of 86 ± 25 from cell therapy. Functional analysis was performed using RV echocardiographic standard and strain parameters in all animals at each step of protocol: at baseline before rTOF surgery, at 5 months before cell therapy and at 7 months at the end of study.

### Echocardiography

Echocardiography was performed on closed-chest animals under general anesthesia in dorsal decubitus. We used commercially available Philips Epiq 7 ultrasound system with X5-1 matrix array transducer (Philips Healthcare) or a VIVID E9 (General Electric Medical System, Milwaukee, USA) equipped with a 2.5 Mhz (1.5-4.5) transducer. The values of all echocardiographic parameters were obtained as the average value of three consecutive cardiac cycles during transient apnoea. Measures were indexed to biological length (BL) or body surface area (BSA), calculated using the formula*(53)*

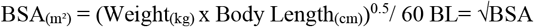

Echocardiographic measurements were determined as previously described *(54)*.

In the parasternal long axis view, the RV anterior wall thickness and the ratio of RV/LV diastolic diameters were measured using TM mode. In the apical four-chamber view, RV end-diastolic and RV end-systolic areas were measured. This allowed the calculation of RV fractional area change (FAC), defined as followed:

FAC (%) = ([RV end-diastolic area – RV end-systolic area]/RV end-diastolic area)*100. Tricuspid Annular Plane Systolic Excursion (TAPSE) was measured as the maximal excursion of the lateral annulus in the apical four-chamber view using M-mode recording through the apex of the RV and the junction between the RV free wall and the tricuspid annulus. Peak systolic velocity of S’wave (Peak S’) was measured at the lateral tricuspid annulus using pulsed tissue Doppler imaging. From a parasternal short axis view, pulmonary annulus diameter and transpulmonary gradient through the pulmonary band were both measured by continuous-wave Doppler flow at the maximal pulmonary valve opening during systole. The presence of a diastolic flow reversal of color Doppler flow in the main or the branch pulmonary arteries was used to predict severe pulmonary regurgitation.

Among the echocardiographic measures, RV end-diastolic and end-systolic area, and RV/LV ratio reflected RV dilation whereas the mean trans-pulmonary gradients were used to reflect the RV pressure overload. Finally, FAC, TAPSE and Peak S’ represented overall RV systolic function.

Regarding the speckle tracking analysis, RV longitudinal deformation was measured by standard 2D grey-scale acquisitions. The peak longitudinal strain (LS) values were computed automatically, generating regional data from six segments of RV and an average value for apical 4-chamber. The peak LS was also assessed by a semi-automatic contouring of the endocardium of the free wall (FW), excluding the interventricular septum to minimize the influence of the interventricular dependence. The peak global RV FW strain was defined as the average of the three FW segments. Peak LS is a negative value, but for ease of interpretation in comparing serial LS values, we have expressed LS as an absolute value. All images were recorded at a frame rate of 50-80 frames per second were stored for postprocessing analysis with QLAB 10 CMQ (Philips) advanced quantification software (EchoPAQ, 110.1.2 GE Healthcare, Horten, Norway)

Clinical status was evaluated daily during the study. Histologic analyses were performed on each animal at the end of the procedure.

### Histology, Immunofluorescence Antibodies

Histology and Indirect immunofluorescence were performed on free wall and HD-area frozen sections. Fibrosis was stained by Sirius red (Sigma-Aldrich, France).

Immunofluorescence used anti–human nuclear KU80 antigen (Cell signaling Technology, C48E7) or anti human mitochondria antibody (anti-CoxIV, abcam ab197491) to detect human progenitor cells, anti-NKX2.5 (R&D AF2444), anti-sarcomeric actinin (Sigma-Aldrich clone E453), anti-NFκB (Cell Signaling Technology, 8242), anti OCT4 (Santa-Cruz Biotechnology rabbit anti-Oct4). Anti-Iba1 antibody (GTX89792) was from GeneTex.

Genomic DNA was prepared from RV free wall and HD-area pieces using the Wizard® Genomic DNA Purification Kit (Promega, France). Samples were submitted to polymerase chain reaction analysis, to quantify the presence of human cells using amplification of human ALU sequences. The negative control DNA was isolated from sham RV pieces.

### Isolation of Pig right ventricular cells

Pigs were euthanized by injection of sodium pentobarbital (10 mL from 200 mg/mL stock) and their hearts were rapidly excised. The aorta was cannulated and rinsed with an ice-cold cardioplegic solution containing (in mM): 110 NaCl, 1.2 CaCl2, 16 KCl, 16 MgCl2, 10 NaHCO3, 9 Glucose supplemented with Heparin (2500 UI/L). The RV was dissected and perfused through the right coronary artery with cold cardioplegic solution while arterial leaks were sutured. The isolated RV was then installed on a Langendorff apparatus and perfused (20 mL/min) with an isolation solution containing (in mM) 130 NaCl, 5.4 KCl, 0.4 NaH2PO4, 1.4 MgCl2·6H2O, 5 HEPES, 10 glucose, 20 taurine, 10 creatine (pH adjusted to 7.4 with NaOH), and supplemented with 0.1 mM EGTA. After this, EGTA was replaced by collagenases (type I, Worthington, USA) and proteases (type XIV, Sigma-Aldrich). Small pieces of myocardium were then dissected and agitated at 37°C in the enzyme-containing solution and myocytes regularly collected by filtration through a 200µm nylon mesh. Cell suspension was then filtered and myocytes were resuspended in Ca^2+^ free Hanks solution and frozen.

### Single–cell RNA-seq

The VD cells were thawed out in PBS EDTA 1 mM and 0.5% BSA. Cells were processed using the Asteria SCIPIO Bioscience kit according to the manufacturer. The cDNA was amplified within a library (NEB kit) and sequenced. The quality control, clustering and further analysis was performed using SCHNAPPS *(55)*

### Spatial transcriptomics

FISH with rolling circle amplification (RCA) was performed using the Xenium (10X Genomics) device. A Xenium Advanced Panel of 218 Sus scrofa genes (V1 chemistry) was designed with Panel designer (version 3.4.7) to identify heart cell types with the top gene markers from the nucseq experiments (Design number 7HVPAE, 10X Genomics)*(56, 57)*. A total of 6 fresh frozen 10 µm pig heart samples were cryo-sectionned using a Leica CM3050S cryostat, then placed on 2 Xenium V1 slides (10X Genomics, #174825). Processing was performed according to the manufacturer’s user guides (CG000581_RevD, CG000749_ RevB, CG000584_ RevK). Gene panel hybridization was performed for 22h, followed by imaging, then quantification, according to the vendor guidelines (instrument software, version 3.2.1.2; analysis version, xenium-3.2.0.7). After a DAPI staining, nuclear segmentation was done, and a cell by gene table count was generated. At the end of the Xenium experiment, sections were colored by an hematoxylin-eosin staining. Gene table count was transformed with spatial data-io (Version 0.1.7) into a SpatialData object that incorporated cell metadata and cell spatial coordinates*(57)*. The object was finally updated to provide a nuclear quantification with sopa (v.2.0.3)*(56)*, in order to directly compare it with the single nucleus data.

### Real time PCR

RNA was extracted from samples with a Quick RNA miniprep kit (Zymo Research, Ozyme, France). QPCR was performed using the LightCycler® FastStart DNA Master PLUS SYBR Green I kit and a light cycler 2.0 (Roche Life Science, France) as previously reported *(16)*

The primers used were:

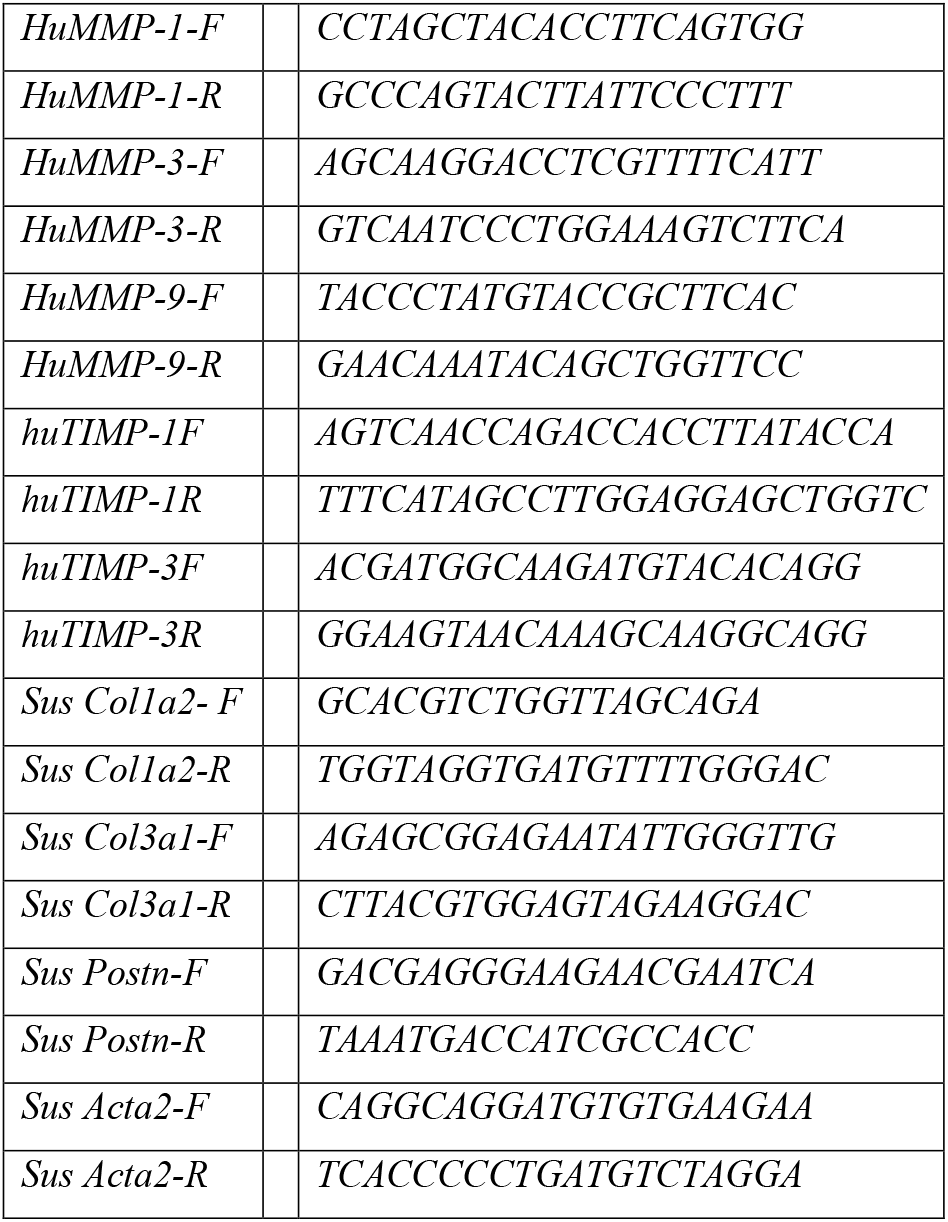
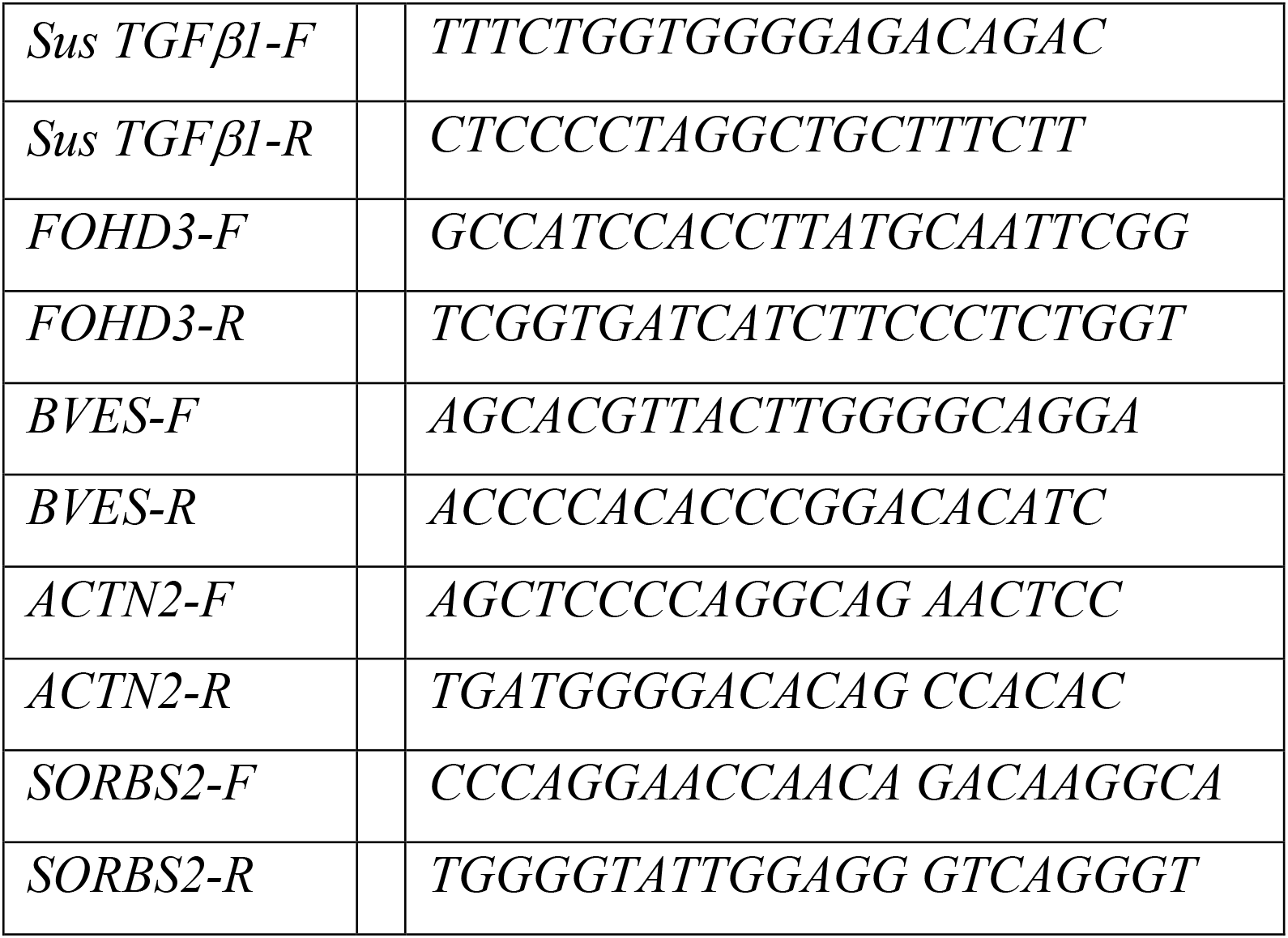

