## Supplementary material for "Macrophage-mediated epimorphosis in pig heart with right ventricular failure": supplemantal figures

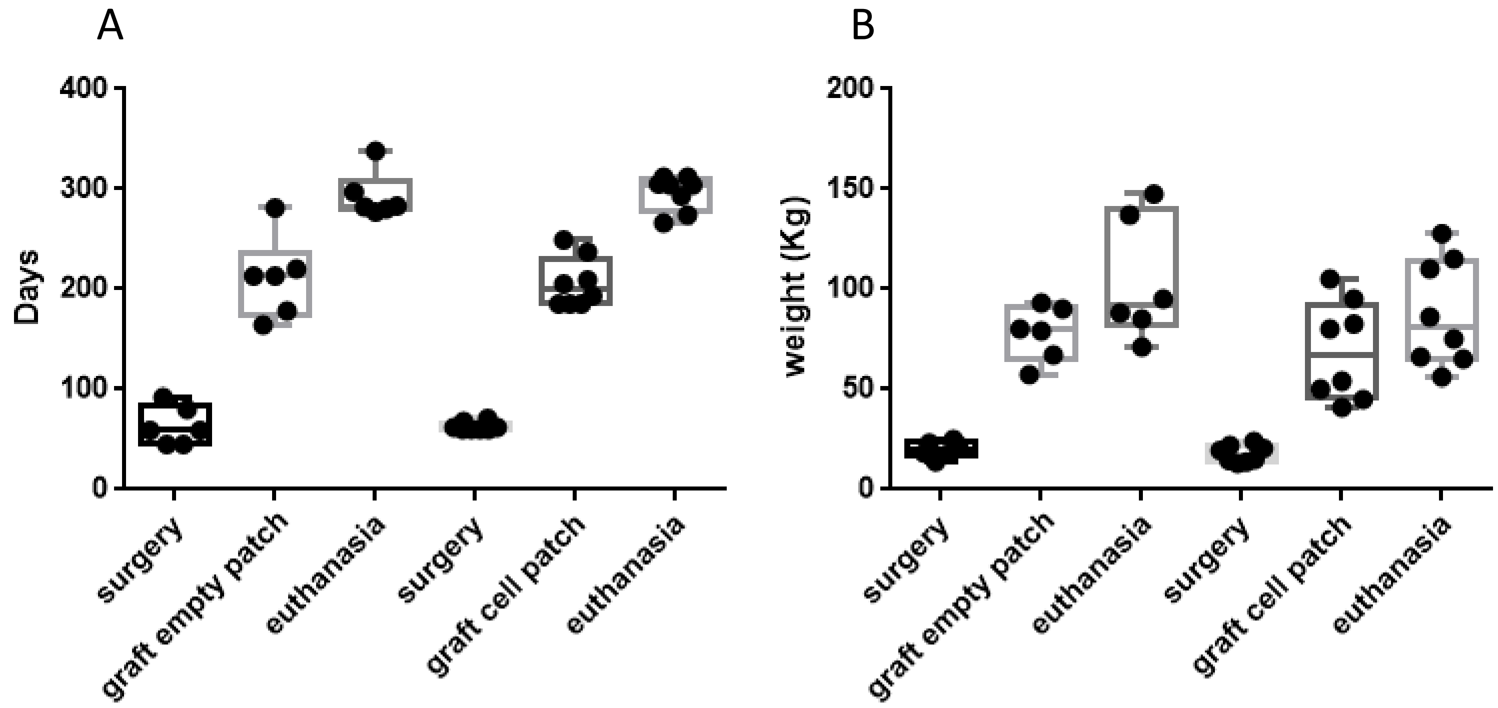


**Figure S1: Pig age (A) and weight (B) in the course of experiment**

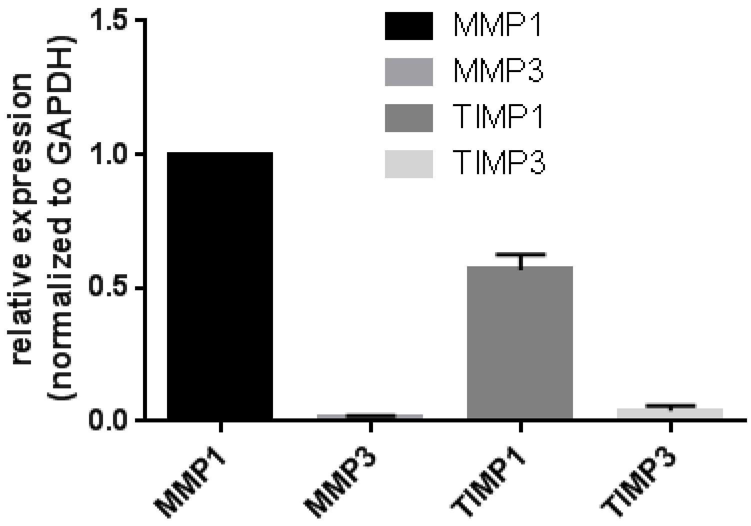


**Figure S2: QPCR of MMP genes from RNA extracted from human ES cells derived cardiac progenitor cells**

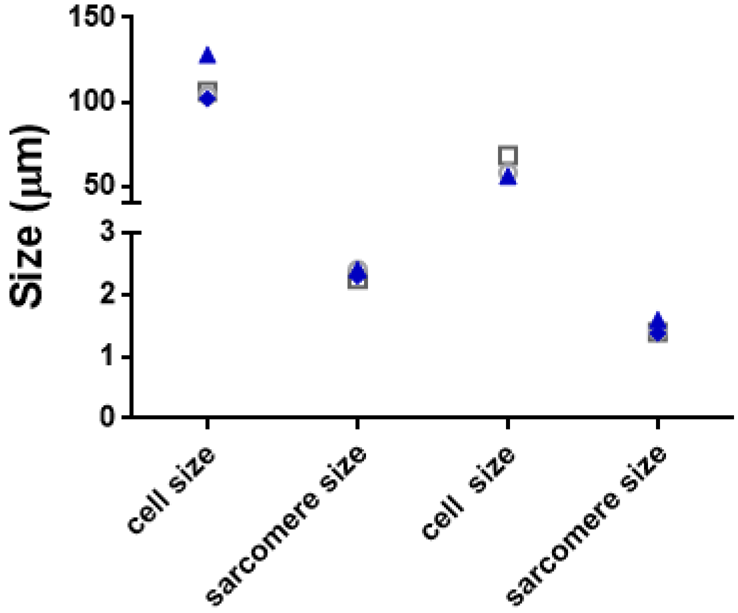


Figure S3 : cell size and sarcomeres in pig RV   (n=72)

A

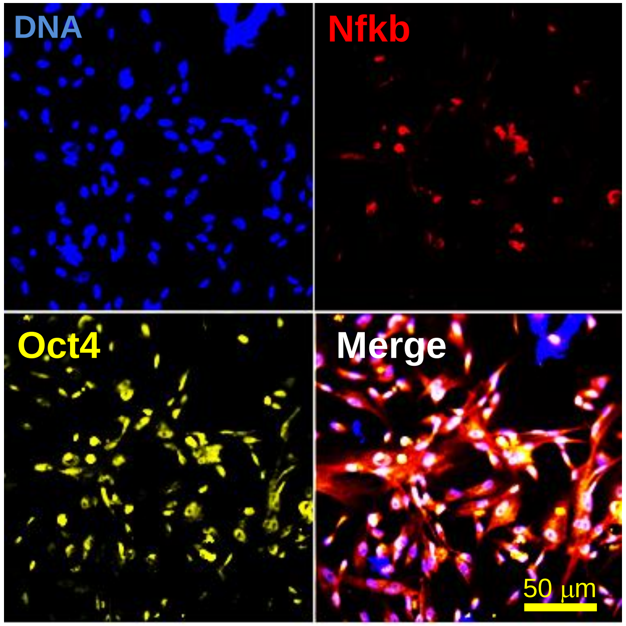


B

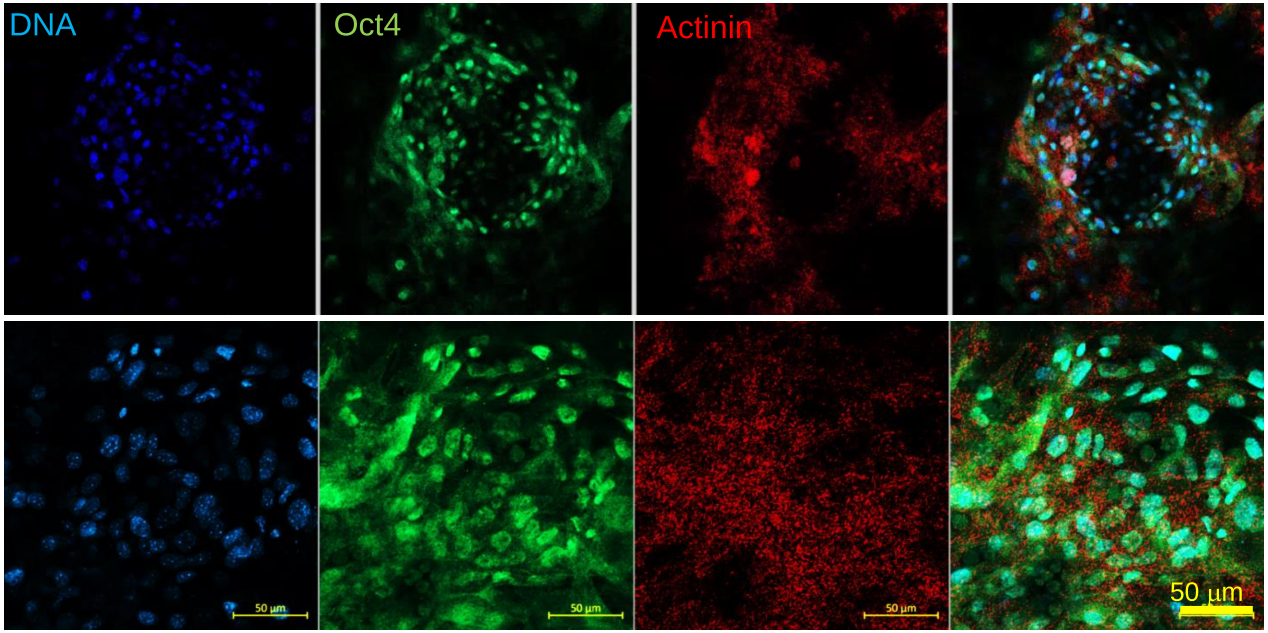


**Figure S4: Nfkb triggers reprogramming of cardiac fibroblasts and cardiomyocytes**. Nfkb (*relAp65*) was transfected in human cardiac fibroblasts (**A**) and mouse postmitotic cardiomyocytes (**B**) . Cells were left in culture for two weeks and stained for Oct4 and actinin. The experiments were performed in triplicate

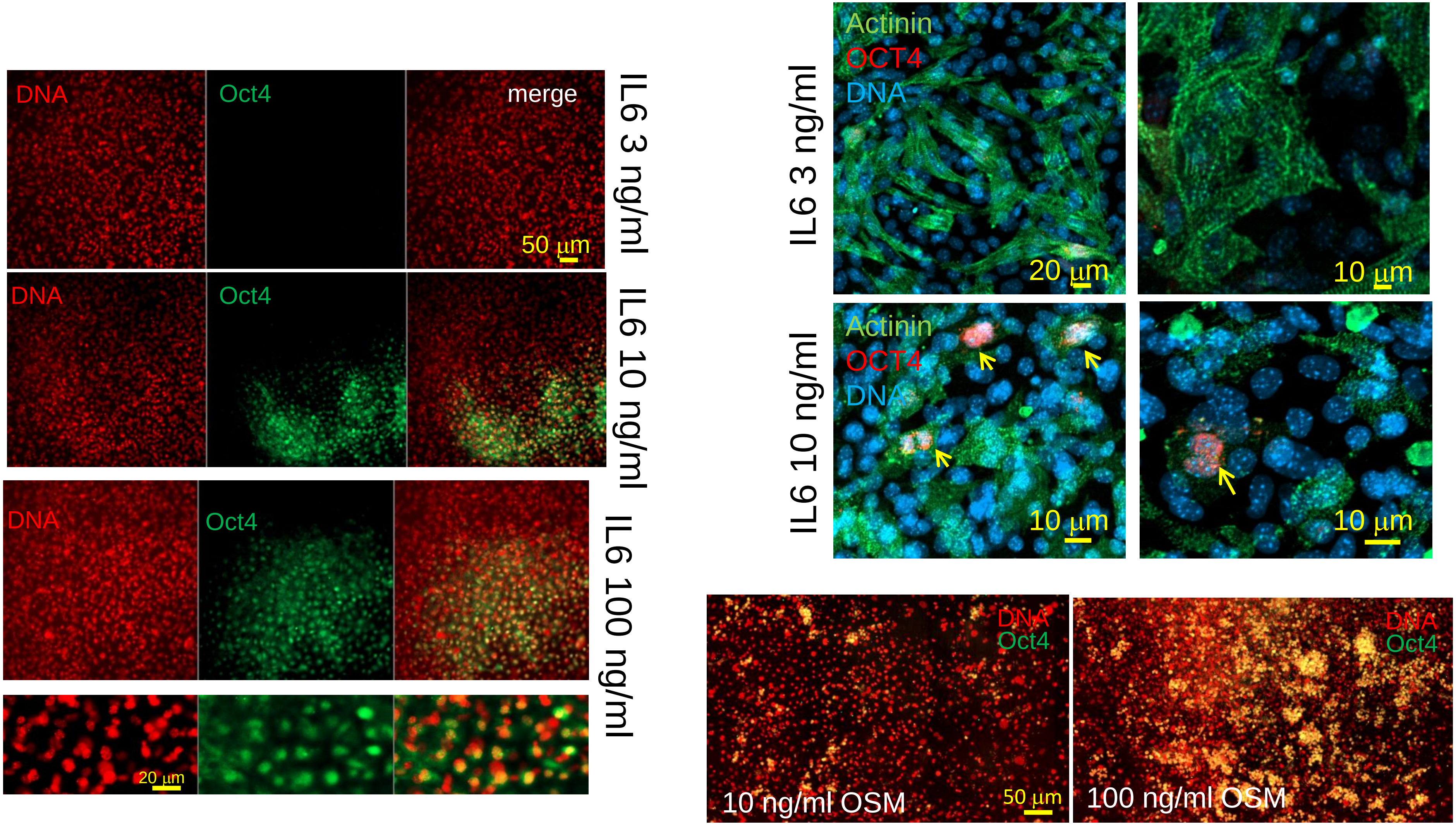


**Figure S5: IL6 and oncostatin reprogram 7 days old post-mitotic mouse cardiomyocytes in a dose dependent manner.** Cardiomyocytes in culture were treated for 7 days with 3, 10 or 100 ng/ml IL6 , or with 10 and 100 ng oncostatin-M then stained with anti-Oct4 antibody and dapi (red). Cells were counterstained with anti-actinin antibody. Yellow arrows point to OCT4+ nuclei.
